# Hippocampal spatial memory representations in mice are heterogeneously stable

**DOI:** 10.1101/843037

**Authors:** Samuel J Levy, Nathaniel R Kinsky, William Mau, David W Sullivan, Michael E Hasselmo

## Abstract

The population of hippocampal neurons actively coding space continually changes across days as mice repeatedly perform tasks. Many hippocampal place cells become inactive while other previously silent neurons become active, challenging the belief that stable behaviors and memory representations are supported by stable patterns of neural activity. Active cell replacement may disambiguate unique episodes that contain overlapping memory cues, and could contribute to reorganization of memory representations. How active cell replacement affects the evolution of representations of different behaviors within a single task is unknown. We trained mice to perform a Delayed Non-Match to Place (DNMP) task over multiple weeks, and performed calcium imaging in area CA1 of the dorsal hippocampus using head-mounted miniature microscopes. Cells active on the central stem of the maze “split” their calcium activity according to the animal’s upcoming turn direction (left or right), the current task phase (study or test), or both task dimensions, even while spatial cues remained unchanged. We found that different splitter neuron populations were replaced at unequal rates, resulting in an increasing number of cells modulated by turn direction and a decreasing number of cells with combined modulation by both turn direction and task phase. Despite continual reorganization, the ensemble code stably segregated these task dimensions. These results show that hippocampal memories can heterogeneously reorganize even while behavior is unchanging.

**Significance statement:** Single photon calcium imaging using head-mounted miniature microscopes in freely moving animals, has enabled researchers to measure the long term stability of hippocampal pyramidal cells during repeated behaviors. Previous studies have demonstrated instability of neural circuit components including dendritic spines and axonal boutons. It is now known that single units in the neuronal population exhibiting behaviorally relevant activity eventually become inactive and that previously silent neurons can quickly acquire task-relevant activity. The function of such population dynamics is unknown. We show here that population dynamics differ for cells coding distinct task dimensions, suggesting such dynamics are part of a mechanism for latent memory reorganization. These results add to a growing body of work showing that maintenance of episodic memory is an ongoing and dynamic process.

## Introduction

The belief that stable behaviors and reliable memory representations are supported by stable elements of neural circuits (Barnes et al., 1997; Thompson & Best, 1990) has been challenged by many findings that neural circuit components across the brain are unstable over time. Circuit instability is notable in the continual replacement of active cells with previously silent cells (Kinsky et al., 2018; Mau et al., 2018; Ziv et al., 2013), but is also observed in the impermanence of dendritic spines and axonal boutons (Attardo et al. 2015; Pfeiffer et al. 2018; Grutzendler et al. 2002; De Paola et al. 2006). How circuit instability may affect neural function is a topic of much debate (Chambers & Rumpel, 2017; Rule et al., 2019).

In the hippocampus, a hub for episodic memory and spatial navigation, change is observed in the patterns neuronal of activity and the set of currently active cells. In behaving animals, single neurons become more sensitive to task demands during training and change their firing properties to more precisely encode task demands (Kobayashi et al. 2003; Komorowski et al. 2009; Lever et al. 2002). Hippocampal memory representations are also unstable even during over-trained behaviors, exhibiting a decorrelation in ensemble activity relative to the elapsed time between recordings (Mankin et al. 2015; Mankin et al., 2012; Rubin et al. 2015; Ziv et al., 2013). These decorrelations result both from remapping of firing locations exhibited by continuously active single neurons that is unrelated to changes in behavior (Mehta et al. 2000; Poe et al. 2000; Lee et al. 2006; Law et al. 2016), and from population dynamics that include the continual inactivation of active cells and their replacement by previously silent cells (Mau et al., 2018; Ziv et al., 2013). However, these changes have primarily been observed during learning or during performance of foraging tasks. How changes occur during stable performance of a multi-dimensional memory task remains an open question. Previous studies have linked the long term stability of a neuronal activity to different spatial locations and different task behaviors (Kentros, et al., 2004; Kinsky et al., 2019; Taxidis et al., 2018). We sought to expand on these studies by examining how different demands on long term memory influence the evolution of hippocampal memory representations during a task where mice pass through the same spatial location under multiple different task conditions.

To study the reorganization of hippocampal representations over time, we used *in vivo* calcium imaging to monitor the activity of hundreds of neurons across multiple sessions in mice performing a Delayed Non-Match to Place task on a figure-eight maze. We first confirmed that neurons modulate their activity on the central stem according to the animal’s upcoming turn direction and the current task phase (Griffin et al., 2007; Wood et al., 2000). We show that the distribution of these single unit responses among the active population changes over time, resulting in an increased number of turn direction-modulated neurons and a decrease in the number of neurons modulated by both the current task phase and upcoming turn direction. These changes primarily result from the unequal recruitment of previously inactive cells to different neuron coding types. While the distribution of single unit activity was unstable, population analyses revealed a stable separation of task variables in the collective ensemble at extended lags between recordings. These results demonstrate that behavior and population output can remain stable while single neuron responses are unevenly reorganized.

## Methods

### Surgical Procedures

4 male, naïve mice (C57BL6, Jackson Laboratory) underwent two stereotaxic surgeries to prepare for calcium imaging. All procedures presented here were approved by the Institutional Animal Care and Use Committee (IACUC) at Boston University. Mice were given 0.05mL/kg buprenorphine as a pre-surgical analgesic, and were anesthetized with ∼1% isofluorane delivered with oxygen. The first surgery was to infuse virus to express GCaMP6f. A small craniotomy was made above the dorsal hippocampus at AP −2.0mm, ML +1.5mm relative to bregma, and the infusion needle was lowered at this site to DV −1.5mm. 350 nL of the viral vector AAV9-Stn-GCaMP6f (University of Pennsylvania Vector Core, obtained at a titer of ∼4×10e13GC/mL and diluted it to ∼5-6×10e12GC/mL with 0.05M phosphate buffered saline) was infused at 40nL/min and allowed to diffuse for 15 minutes before the infusion needle was slowly removed.

The second surgery, to implant a gradient-index (GRIN) lens for imaging, was performed three weeks later to allow for viral infection and GCaMP6f expression. A 2mm diameter circular craniotomy was made at AP-2.25mm, ML +1.8mm, and the neocortex was aspirated until rostral-caudal fiber tracts of the alveus were visible. Near-freezing 0.9% saline solution and GelFoam (Pfizer) were used continuously to control bleeding and to dry the base of the craniotomy prior to lens implantation. The GRIN lens (1mm diameter, 4mm length, Inscopix) was slowly lowered stereotaxically to 200 um dorsal to the infusion site of the virus, measured relative to the skull surface. The lens was then fixed in place using a non-bioreactive silicone polymer (Kwik-Sil, World Precision Instruments) to entirely cover the craniotomy, which was then covered with Metabond dental cement (Parkell) to anchor the lens to the skull. The lens was covered with a temporary cap made from Kwik-Cast (World Precision Instruments) until the baseplate was attached.

After allowing a week of recovery from the lens implantation surgery, mice were again anesthetized and placed in the stereotaxic holder. The baseplate was magnetically attached to the imaging microscope camera, which was then aligned parallel to the GRIN lens by adjusting until the edge of the lens was entirely in focus in the nVista recording software (Inscopix). The camera with baseplate was then lowered until GCaMP6f-expressing cells were optimally in focus, and then raised by 50 um to allow for shrinkage of the dental cement used to affix the baseplate. The baseplate was then fixed in place to the existing metabond around the GRIN lens with Flow-It ALC Flowable Composite (Pentron), and cured with ultraviolet light. Gaps in the dental cement were filled in with Metabond, the camera was removed, and a cover attached to the baseplate.

### Maze Description

The maze was constructed from wood and the internal floor area measured 64.5 cm long by 29.2 cm wide, and walls were 17.75 cm high. Middle maze walls separated this area into a central hallway (Center Stem) and left and right Return Arms. Each hallway was 7.5 cm wide. This resulted in low variability of the animals’ left/right position within a hallway, although it did not prevent the animals from occasionally running with their head turned towards one side. Rewards were delivered through ports at the maze walls at floor level of the side arms 12 cm from the delay-end of the maze. To dictate turn direction on Study Trials (see below) and to contain the mouse during the delay period, arm barriers were used that were made of transparent plastic. The delay barrier was made of wood. In this manuscript we only consider data from the central stem and return arms.

For analysis of the central stem, we chose a region starting ∼8 cm in front of the delay barrier and extending 30cm to end ∼5 cm before the choice region at the end of the middle maze walls; this region was selected to encompass the region where the mouse was running similarly between study and test task phases and left and right turn directions. Left and right variability in the animals’ head position at the end of this region was less than 2.5 times the standard deviation of the animals’ left/right variability for the first half of the stem, and was usually indistinguishable by visual observation in behavioral recordings. We divided this 30cm long region into 8 spatial bins each 3.75 cm in length. For the return arms (**Supplement**), we chose a region of equal length that started after the animals had fully entered the return arms and ended before they reached the reward zone, also 30cm in length and separated into 8 bins each 3.75 cm.

### Behavior pre-training and recording sequence

Mice were trained to run on a Delayed Non-Match to Place (DNMP) task shown in **Figure 1**. This involved extensive pre-training in order to obtain performance at the criterion of 70% correct.

**Figure 1.**
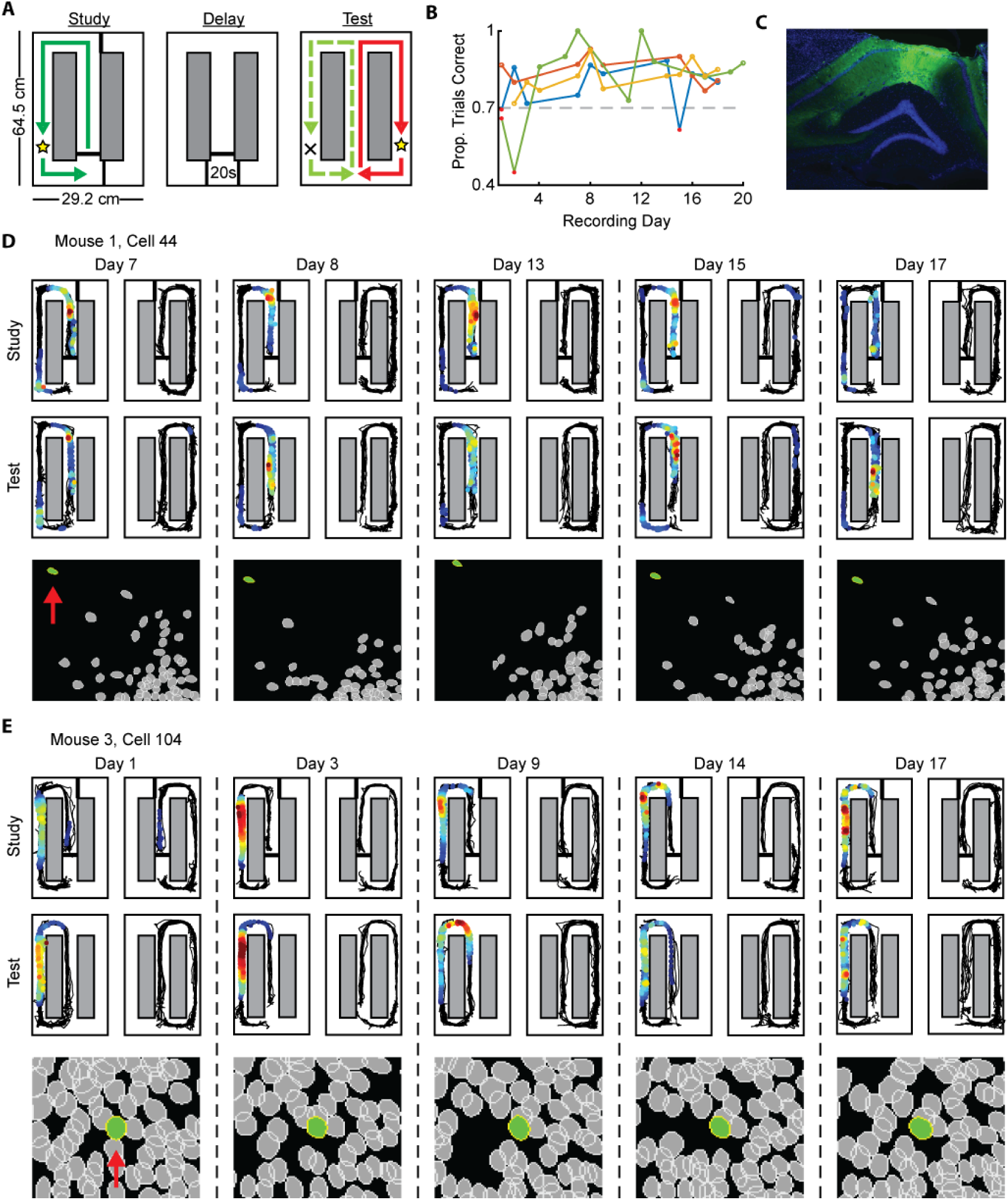
***A,*** Task outline: each trial has a Study and Test Phase, separated by a 20-second delay. Each trial is followed by a 15-25s inter-trial interval in the mouse’s home cage, adjacent to the alternation maze (not shown). ***B,*** Performance of individual mice (separate colors) over all days of recording. Only sessions with performance above 70% were included, excluded sessions are marked in red. ***C,*** Example viral expression and lens placement in dorsal CA1. Green is GCaMP6f-EYFP, blue is DAPI. ***D,*** Top: Activity maps for one cell (a turn splitter neuron; see Figure 2) over five days of recording. Each plot represents the average activity map for one task condition combination, ordered clockwise from top-left: Study-Left, Study-Right, Test-Right, Test-Left. In each plot, the black trace is the animal’s recorded position, and colored dots indicate frames where the cell was active. Dots are colored based on the local event likelihood, normalized by local occupancy, where red is the highest event likelihood within that day and blue is the lowest. Bottom: Cell ROI masks for that recording day. Cell of interest is colored in green, and indicated with red arrow on first day shown. Masks were aligned across days based on relative positions of cells and cells were aligned based on the distance between cell centers and correlation of masks (see Methods). ***E,*** Same as **D** but for a cell with an activity field on one return arm.

After fully recovering from surgeries, mice were extensively handled for ∼15 min/day for 5 days. They were simultaneously food restricted to 80% of free feeding body weight, and acclimated to consuming chocolate sprinkles. Over the next two weeks, mice were given time to explore the maze, and were slowly shaped to run in a single direction through the maze and to receive reward, with inserted walls to block paths and guide them. In the last few days of pre-training, mice were guided with blocking walls to alternate between the two reward arms and given experience with continuous and delayed alternation.

Mice were recorded performing two tasks. In the Delayed Non-Match to Place (DNMP) task (Griffin et al., 2007), mice alternated between Study and Test trials. On Study trials, mice were placed in the center stem in front of the delay barrier, ran to the choice point, where a removable barrier forced them to take a path down one return arm where they received a reward of one chocolate sprinkle. They then moved to the delay area, waited through a 20-second delay, and the delay barrier was lifted to start the Test trial. On a test trial, mice again ran to the choice point but there was no barrier and mice had to go down the return arm opposite to the preceding study trial in order to receive a reward. They then moved to the delay area, from which they were removed to their home cage to wait through a 15-25 second inter-trial interval while the next Study trial was prepared. Mice completed between 25 and 40 Study-Test trial pairs per session.

A second task, termed the Forced-Free task, was used on other days for a different study question not addressed here. On each trial in the Forced-Free task, mice were placed in front of the delay barrier, proceeded to the choice point and were either forced down a particular return arm or were free to choose which arm. On all trials mice received a reward regardless of which arm they entered. After consuming the reward, mice entered the delay area and were immediately returned to their home cage for a 15-25 second inter-trial interval while the next trial was prepared. Mice typically completed 40 trials per session. Forced and free trials were pseudo-randomly interleaved, as was turn direction on forced trials.

The full recording sequence was two rounds of the following sequence: one day of Forced-Free, 3 days of DNMP, and one day of Forced-Free. This was followed by a sequence with one day of Forced-Free followed by 5 days of DNMP, followed by one day of Forced-Free. Gaps between Forced-Free-DNMP recording sequences ranged between 0 and 2 days (Full sequence: FF-D-D-D-FF, break, FF-D-D-D-FF, break, FF-D-D-D-D-D-FF). Data from the Forced-Free task are not presented here.

We only include data from DNMP recordings where cell registration could be reasonably performed and where the animal’s performance was ≥70%.

### Imaging

Imaging data were acquired using a commercially available miniaturized head-mounted epifluorescence microscope (Inscopix). Microscopes were attached on awake, restrained mice, and optical focus, LED gain and intensity adjusted for each individual mouse but kept stable across days. Videos were captured at 20 Hz with a resolution of 1440 x 1080 pixels, spatially downsampled 2x to 720 x 540 pixels. Dropped and corrupted frames were replaced with the preceding good frame, and lost frames were excluded from analysis. Mosaic (Inscopix) was used to pre-process recordings for motion correction and cropping (exclude pixels without GCaMP6f activity), and to generate a minimum projection of the final video (image which has the same height and width of each frame and each pixel is the minimum of that pixel for the entire video) to be used during ROI extraction.

To extract neuron regions of interest (ROIs) and calcium event times, pre-processed videos were then passed through custom-made MATLAB-based image segmentation software (Mau et al., 2018; Kinsky et al., 2018) (TENASPIS, software available at https://github.com/SharpWave/TENASPIS; see D.W. Sullivan et al., 2017, Soc. Neurosci., abstract). Briefly, TENASPIS applies an adaptive thresholding process on a frame-by-frame basis to a band-pass filtered video to identify discrete regions of fluorescent activity (blobs). Blobs are then identified as likely cells based on expected shape and size, and the software aligns these blobs together over successive frames. Dynamics in calcium activity, including event duration, distance traveled over successive frames, and probable spatial origin, are used to identify putative neuron ROIs. Fluorescence of neuron ROIs is refined into events based on the rising phase of calcium activity. Finally, neuron ROIs with significant spatial overlap and high correlations in calcium activity are merged into single cells.

Cells were registered across sessions using a semi-automated procedure with custom software developed in MATLAB that is available along with the rest of our analysis code. For each animal, each session was first aligned to the same ‘base’ session, selected from the middle of the recording schedule. To align sessions, a set of 25-40 ‘Anchor’ cells was chosen based on the relative positions of neuron ROIs in the base session and each other session (**Supplementary Figure 1a-b**). Centers of these ‘anchor’ cells were used to compute an affine geometric transformation (‘fitgeotrans’ function in MATLAB) and then align the entire set of ROIs in the sessions being registered with the base session (‘transformpointsforward’ function in MATLAB). Cells with centers within 3um (translated to pixels) were identified as the same cell, and when there was more than one match within that radius, the registered cell with the higher spatial correlation to the base cell was chosen (**Supplementary Figure 1c**). Cells from a registered session that were not partnered to the base session were added to the set of unique footprints alongside base session cells so that cells in successively registered sessions could be paired to them in turn. Alignment maps were validated by visual inspection: this included looking at the relative alignment with other cells in the field of view, and orientation of putatively mapped cells across sessions. Cells that were not aligned by the automated procedure based on center-to-center distance but that shared orientation and relative alignment to neighboring cells were registered manually (**Supplementary Figure 1e**, green cell). When looking at the relationship for all cell pairs across all sessions, the correlation of ROIs and distances between centers formed a cluster near the top of the distribution for all cell pairs (**Supplementary Figure 1d**). The TENASPIS algorithm is designed to discriminate between partially overlapping cells, which gives rise to in many pairs of cells that have high ROI correlations and low center-to-center distances, but remain unregistered because a better matched pair was found using the procedures above; in **Supplementary Figure 1d**, this manifests in the black points mixed in among the red registered cell pairs.

### Behavioral Tracking

Animal position was recorded using an overhead video camera and CinePlex V2 tracking software (Plexon). Tracking was performed at 30 Hz, and was synchronized with a TTL pulse to the imaging data acquisition through nVista software. Tracking was validated manually and errors were corrected using custom software written in MATLAB. Position was then interpolated to the 20 Hz imaging time stamps.

### Histology

Mice were perfused transcardially with 10% phosphate buffered saline until outflow ran clear and then with 10% phosphate buffered formalin. Brains were then extracted and post-fixed in formalin for 2-4 days, and then transferred to 30% sucrose solution in phosphate buffered saline for 1-2 days. Brains were then frozen and sliced into 40 um sections on a cryostat (Leica CM 3050S), mounted, and coverslipped with Vectashield Hardset mounting medium with DAPI (Vector Laboratories). Slides were then imaged using a Nikon Eclipse Ni-E epifluorescence microscope at 10x and 20x to verify viral expression and location and GRIN lens location relative to the CA1 cell layer.

### Quantification and Statistical Analysis

#### Event likelihood

Calcium events were detected and analyzed to compute the likelihood of calcium events occurring at a given location. The analysis software, TENASPIS, (see above) defines an event as the time during the rising phase of a spike in calcium fluorescence in a cell which exceeds a local threshold of that cell’s session average of fluorescence activity. This returns a binary output for each cell which describes whether that cell was or was not, at every imaging frame, exhibiting a calcium event. We calculated event likelihood by pooling data from the set of trials of interest for each cell (e.g., Study trials on the stem), and then, for each spatial bin, dividing the number of frames for which an event was occurring by the number of frames when the mouse was in that bin in that set of trials. This produces an output between 0 (an event never occurred in that spatial bin) and 1 (an event always occurred when the mouse was in that spatial bin).

#### Active Cells

For single unit analyses, cells are included on a given day when they exhibited a calcium event on at least 25% of trials or 3 consecutive trials in a single trial type (e.g. Study-Left). In the population analyses, we included all cells were successfully registered to the sessions being compared.

#### Splitter Identification

Splitter neurons are cells that exhibit a significant bias in their firing activity on the central stem for trials of a particular upcoming turn direction (Left versus Right) or task phase (Study versus Test) (**Figure 2**). Thus, each cell is a member of one of four mutually exclusive categories, depending on whether its calcium activity is modulated by either task dimension, both, or neither: turn splitter neuron, task phase splitter neuron, turn*phase splitter neuron, or non-splitter. Note that turn*phase splitter neurons refer to cells splitting both turn direction and task phase.

**Figure 2.**
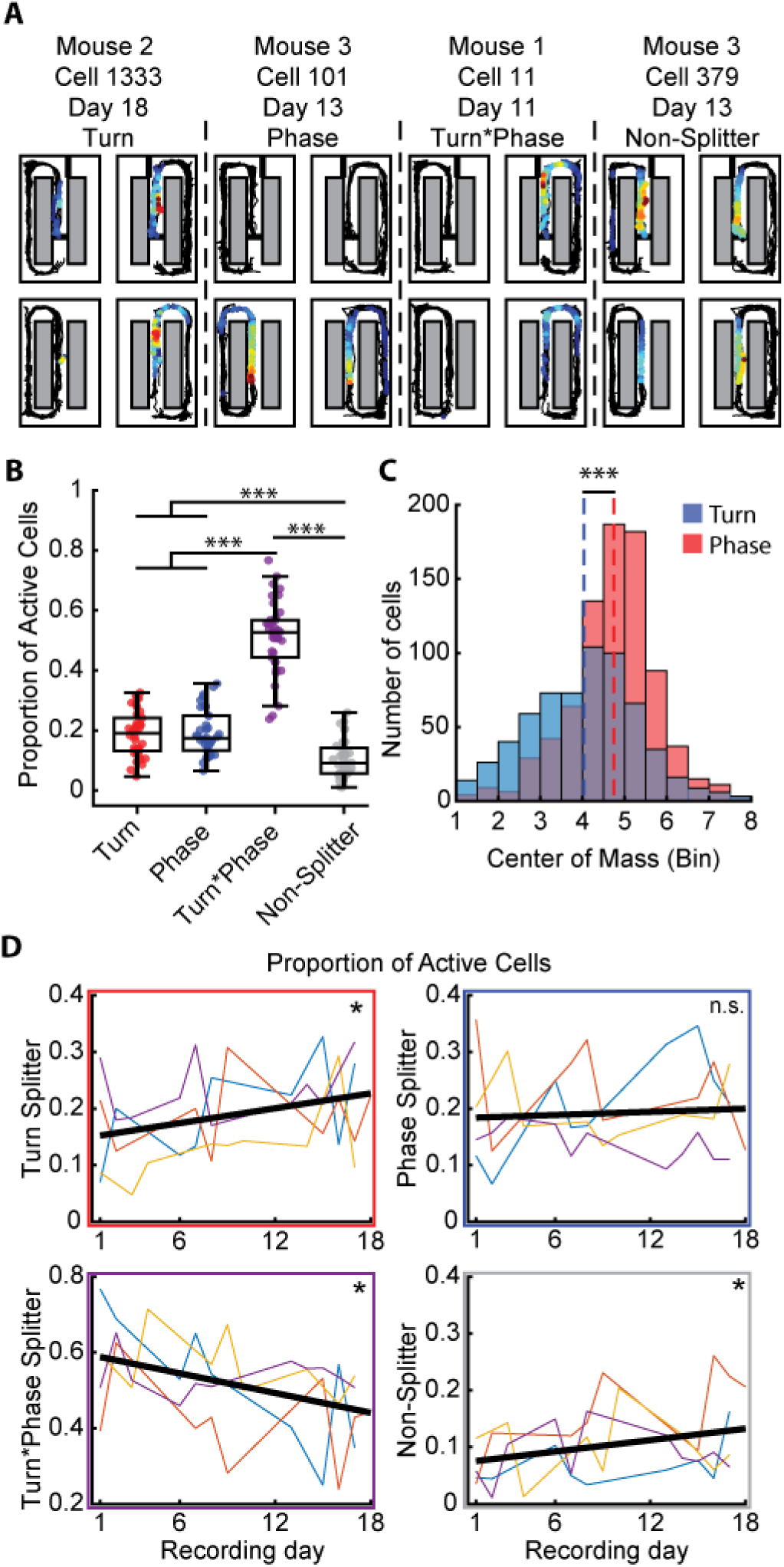
***A,*** Example activity maps for each type of splitter on the central stem. Warmer colors indicate higher transient likelihood. ***B,*** Proportions of splitter cells out of the total active cell population on each day for all animals. Box shows inter-quartile range and middle line shows median. Statistic: Wilcoxon signed-rank test. ***C,*** Distribution of centers-of-mass of event activity for Turn and Phase splitter neurons. Statistic: Mann-Whitney U-test. ***D,*** Proportion of splitter neurons in individual animals (unique colors) and group regression (black) over the course of the experiment. Color of box indicates cell type as described by y-axis label. Significance calculated with Spearman rank correlation between proportion of splitters and recording day number for all included sessions (n=38). * p<0.05, ** p<0.01, ***p<0.001

To identify whether each cell’s activity was significantly modulated by task variables, we used a permutation test to measure the significance of the difference in event activity likelihood against a shuffled distribution. This was repeated separately to measure activity bias for turn direction or task phase. We first separated epochs when the mouse ran through the central stem according to the given task dimension (i.e. left and right turn trials, or study and test trials), and computed the event likelihood (see above) for these sets of trials. Then took the difference in likelihood scores by subtracting the Right trial event likelihood in each spatial bin from that for Left trials, or Test trial from Study. We then repeated this for all 1000 sets of shuffled trials, which were generated by shuffling the trials between trial types accordingly, to get a shuffled difference distribution. Cells were determined to “split” the dimension of interest if their original event likelihood difference was greater than 95% of the shuffle differences in any spatial bin.

In the supplemental data, this procedure was repeated in the same fashion for epochs when the mouse ran down the return arms to measure selectivity for the separate (Right or Left) return arms and for Study and Test task phases while on the return arms.

#### Population Vector Correlations

Population vector correlations were computed in a manner similar to that described by Leutgeb et al. (2005)(**Figure 3a**). We generated three sets of correlations: 1) within-condition: trials of the same type (e.g. Study-Left vs. Study-Left); 2) Left vs Right, and 3) Study vs. Test. First, trials were grouped for the comparison of interest and then each group was split so that within condition comparisons would have the same number of trials as the other two correlations. For a given half-set of trials, we computed the event likelihood in each spatial bin with the method described above. We then took these spatial bin event likelihoods for the set of cells included and computed a Spearman correlation for each spatial bin against the event likelihoods in the same spatial bin for the trials in the different comparisons listed above. For correlations computed across days, we computed all day-pair combinations for each self-comparison and for each comparison between study and test trials and between left and right turn trials, for example between left turn trials on day 1 and right turn trials on day 4. Cells included were those present (successfully registered) on both days for each comparison (Similar results were achieved using several other cell inclusion criteria, data not shown).

**Figure 3.**
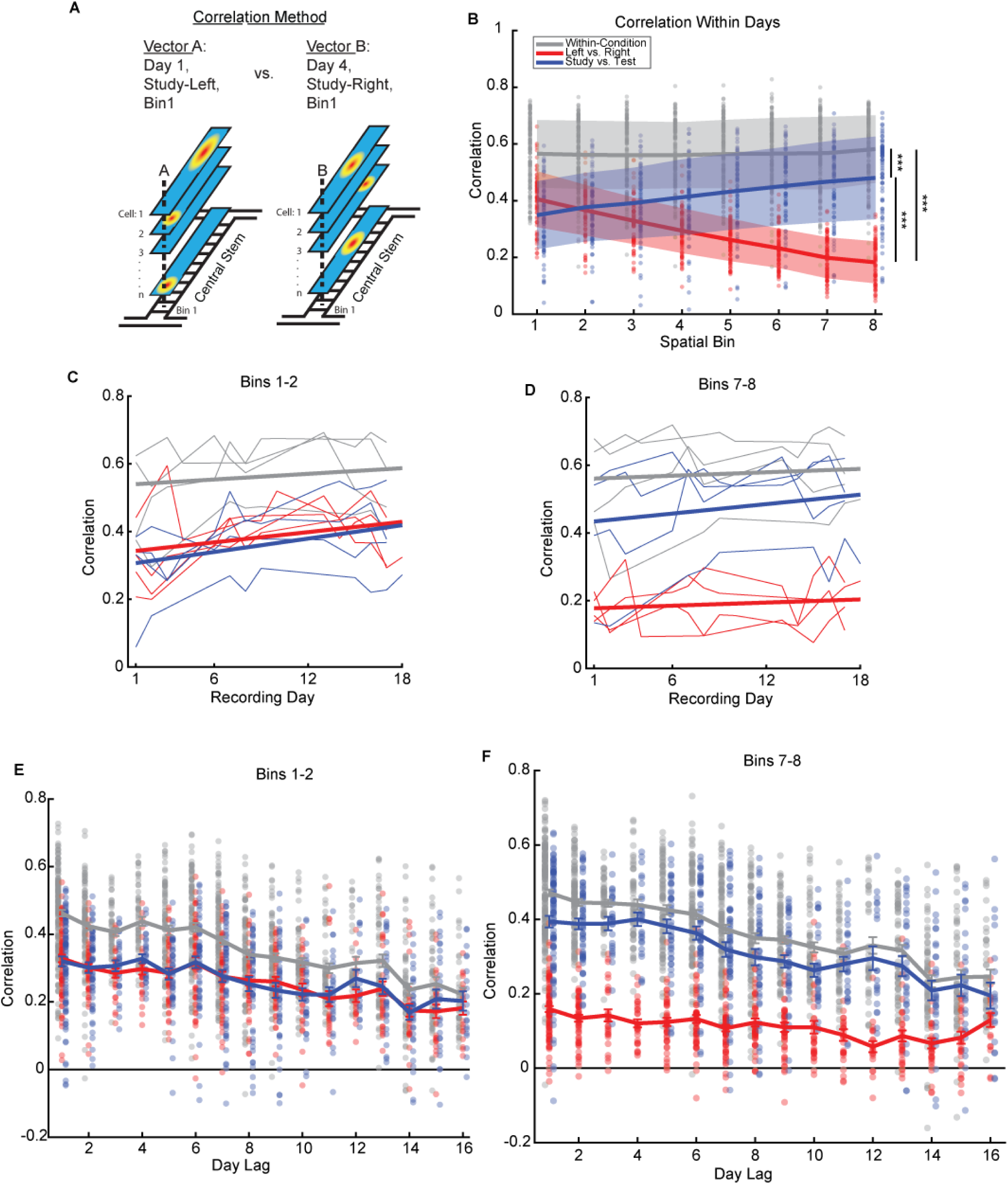
***A,*** Method for making population vector correlations. ***B,*** Population vector correlations between trials of the same turn direction and task phase (gray), different turn directions (red) and different task phases(blue). Correlations in this panel B are generated from trials that occur on the same day. Shaded patch indicates 95% of points for the indicated correlation type in that spatial bin, trend line indicates mean. Statistic: Wilcoxon rank-sum test on all points for these groups. ***C,D,*** Mean correlation for pairs of spatial bins over the course of recordings. Thin lines indicate individual animals’ correlations, bold lines are best fit regression. Statistic: Spearman rank correlation on points from all recording days. ***E,F,*** Correlations between trials on separate recording days for indicated pairs of spatial bins. See text and supplementary data tables for statistics. * p<0.05, ** p<0.01, ***p<0.001

#### Statistics

All statistical tests were done with Spearman rank correlations, Wilcoxon Rank-sum tests (Mann-Whitney U tests), Wilcoxon signed-rank tests, sign tests, or permutation tests with threshold set at >95% of shuffles for the given test. These tests were used because data were often not normally distributed.

## Results

### Heterogeneous changes in daily distribution of single-cell task-related responses

We recorded calcium activity in neurons in dorsal area CA1 as mice performed a delayed non-match to place (DNMP) task over several days. In the DNMP task, mice first run a study trial where they are forced to turn into one side arm to receive reward. After a 20-second delay, mice must choose to go down the opposite arm to receive a reward (**Figure 1a**). We used this task because mice traverse the same section of the maze (the central stem) under each combination of Task Phase and upcoming Turn Direction. This allows us to examine hippocampal representations of the same space under four different behavioral conditions: Study-Left, Study-Right, Test-Left, Test-Right. We recorded 8256 cells in four male mice across 38 sessions with a behavioral performance (opposite turn direction on Test trials relative to preceding Study trial) minimum of 70% (9 days in 3 mice, 11 days in 1) (**Figure 1b**), spanning up to 17 calendar days. Performance did not change over the experiment (only days above threshold: rho=-0.031, p=0.852; all days recorded: rho=0.198, p=0.210; Spearman rank correlation). We recorded activity using the virally-delivered fluorescent calcium indicator GCaMP6f and head-mounted miniature microscopes (**Figure 1c**), and extracted cell ROIs using custom software (example ROIs in **Figure 1d-e**, bottom; see Methods) (Kinsky et al., 2018; Mau et al., 2018). On average, each cell was successfully registered for 3.45 sessions, and cells often displayed stable activity profiles across sessions (**Figure 1d-e**, top).

Single cells often modulate their spatial firing activity according to context-dependent task dimensions such as upcoming turn direction or current task phase. Turn direction responses are thought to represent specific spatial trajectories (Frank et al. 2000; Wood et al. 2000; Ferbinteanu and Shapiro 2003), while a task phase-modulated response profile reflects the (presumably) different network activity states for encoding during the study phase and retrieval during the test phase (Griffin et al. 2007). We assessed whether these task variables were encoded in the calcium activity of neurons in our recordings using a permutation test (see Methods) and found that ∼90% of cells active on the central stem (3443/3810 active on any recording day) displayed a functional phenotype described by a modulation of their calcium activity according to the animal’s upcoming turn direction (turn splitter neurons), the current task phase (phase splitter neurons), or both (turn*phase splitter neurons) (see examples in **Figure 2a**); these categories are mutually exclusive. Note that we found many cells which display a turn direction-modulated response on Study trials, indicating that mice could likely see the turn barrier before having reached it.

On the center stem, there was no difference in the proportions of turn or phase splitter neurons (18.96±1.22% and 19.19±1.20%, respectively, z=0.016, p=0.987, Wilcoxon signed-rank test), but there were more turn*phase splitter neurons than either group (51.44±1.93%, both z=5.286, p=1.250e-07) (**Figure 2b**). We also observed a location bias among different splitting phenotypes of single cells: phase splitter neurons were more likely to have their activity center of mass (event activity pooled across all trial types) closer to the start of the stem than did turn splitter neurons (p=6.719e-30, Mann-Whitney U test) (**Figure 2c**). A bias in firing location may indicate that cells tend to fire in proximity to the behaviors they encode: for phase splitters, this could be whether the trial began in the delay area or being placed on the maze by the experimenter, while turn splitters encode an upcoming spatial turn direction.

The daily distribution of splitter types was not stable: the percentage of turn*phase splitters significantly declined over the course of the experiment (rho=-0.35774, p=0.027, Spearman rank correlation), though it remained greater than other splitter types. Meanwhile, the percentage of phase splitter neurons was stable (rho=0.084, p=0.616) and the percentage of turn splitter neurons went up (rho=0.347, p=0.033) (**Figure 2d**). The percentage of non-splitters displayed a small but statistically significant increase over the course of the experiment (rho=0.331, p=0.043) (**Figure 2d**). The proportions of each type of splitter neuron were not correlated with animals’ performance on the DNMP task (all rho absolute value <0.217, all p>0.190) (**Supplementary Figure 2**). These findings replicate a previous result in a new species (Griffin et al. 2007) and extend that work to show that the distribution of task-dimension modulated responses among neurons is unstable over time, even though behavioral output is reliable. In particular, the number of turn splitter neurons increases over time, whereas the number of turn*phase splitter neurons decreases over time, suggesting representations become less experience-specific over time.

We applied these same analyses to determine neuronal activity modulation according to task variables to neuronal activity during the return arm epochs. Because this analysis is performed in the same way, it can be used to indicate relative distinctiveness in the way neurons code for overlapping spatial trajectories (central stem) as opposed to unique spatial locations (return arms). Many cells displayed a calcium event bias for one arm over the other (place cells, referred to here as “place splitters”), and many cells also showed selectivity for one task phase. The proportions of place and phase splitter neurons on the return arms did not individually change over time, though there was an increase in the number of cells which were active on the return arms but did not show place or task phase selectivity (**Supplementary Figure 3**). These results show that changes in the representation of the task and environment are modulated by memory load, which is low on the return arms and high in the central stem.

In summary, by demonstrating that the distribution of task variable responses among single units is unstable, we show that representations for various task dimensions experienced in the same spatial location and during a similar behavior are heterogeneously stable, with divergent changes based on their coding of the behavioral context.

### Population-level separation of task dimensions is stable over experience

We next asked how these patterns of activity manifested in the activity state of CA1 as a whole. This population analysis was designed to measure the similarity in the pattern of activity among the population of neurons within and across recording sessions. We computed Spearman correlations for the activity in each spatial bin from the start of the stem to the choice point for a given trial type using the calcium event likelihood for each trial type of all cells present in the session pair (**Figure 3a**)(see Methods). We generated three sets of correlations: 1) trials of the same turn direction and task phase (within-condition; e.g. Study-Left vs. Study-Left), 2) trials of different turn directions (Left vs. Right, abbreviated as LvR), and 3) trials of different task phases (Study vs. Test, abbreviated as SvT).

We found a stable ensemble activity pattern when examining the population vector correlations for trials occurring on the same day. Activity states for trials of the same type were significantly more correlated than those both for trials of different direction and trials of different task phase, showing a discrimination in the ensemble-level code for different trial types (see **Supplementary data table 2** for detailed statistics) As shown in **Figure 3b**, the correlations between trials of the same type did not change across spatial bins (rho=0.045, p=0.116; Spearman rank correlation). In contrast, activity states for left and right trials grew more decorrelated as animals approached the choice point (rho=-0.678, p=4.946e-83), and study and test trials were most discriminable at the start of the stem (rho=0.332, p=4.418e-17). The correlation change along the stem follows the center-of-mass distribution for splitter cell firing fields (**Figure 2c**). This pattern of correlations across spatial bins was stable over the course of recordings (all rho absolute value < 0.313, all p > 0.056; Spearman rank correlation of 2-bin mean for each type of population vector correlation value against recording day number) (Examples for bins 1-2 and 7-8 in **Figure 3c-d**). This result demonstrates that, in spite of the changing distribution of single-neuron encoding properties (**Figure 1d**), the population-level distinction between activity states (**Figure 3b**) and its relationship to spatial position is stable over time (**Figure 3c-d**).

We next assessed the correlations within and between trial types for trials on different days. It may be expected that population activity states would diverge with respect to time (i.e., become less correlated) due to cell replacement and changes in the splitter neuron distribution (**Figure 2**). To assess this, we examined the mean population vector correlations at the beginning and end of the stem between sessions recorded 1 to 16 days apart. We observed that all three types of correlations significantly decreased with increasing day lag at both ends of the stem, (**Figure 3e-f**). However, even as correlations decreased, LvR and SvT correlations were significantly lower than those between trials of the same type for at least a week between sessions and in many cases longer (see detailed statistics in **Supplementary Data Table 3,4**).

These results show that constant cell turnover minimally impacts the ability of the population to represent different experiences of the same space over many days of recording and that this representational structure is preserved over time. However, the extent to which the population distinguishes between task dimensions depends on the dimensions being compared, the animals’ physical location, and the temporal lag between experiences.

### Evolution of single-unit to responses is attributable to changing distribution of new cell activity types

We next assessed the origin of the changes in the distribution of splitter neuron types over time. There are several possible sources of change in the splitter neuron distribution: different splitter neuron types could be persistently active for different amounts of time before becoming silent (variable stability); neurons could change their splitter type (splitter type transition); or previously silent neurons could be preferentially allocated to certain splitter types (unequal allocation of newly active cells). We found no evidence of variable stability: cells were equally likely to stay active in later recording days regardless of splitting type (all p>0.05, Wilcoxon rank-sum test between each pair of splitting phenotypes at each day lag) (**Figure 4a**).

**Figure 4.**
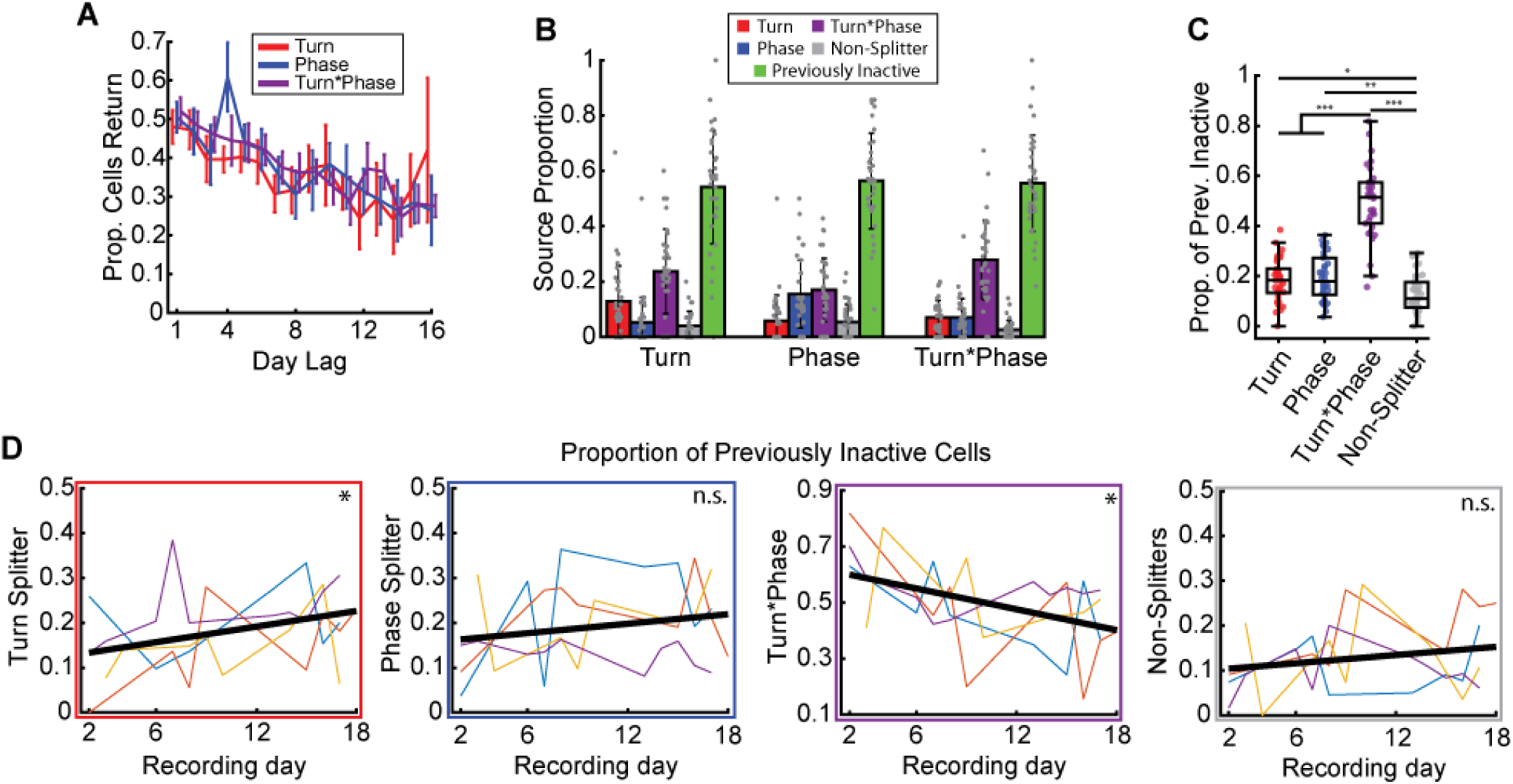
***A,*** Proportion of cells that are still present at increasing day lags. Statistic: Wilcoxon signed-rank test. ***B,*** Proportion of each splitter type by what that cell was on the prior day of recording. ***C,*** Proportion of each splitting phenotype among each recording day’s set of previously inactive cells (from second recording day forward). Statistic: Wilcoxon signed-rank test. ***D,*** Changes in the distribution of splitting phenotypes among previously inactive over the course of recordings. Colored lines are individual animals, black line is best fit regression. Color of box indicates cell type as described by y-axis label. Statistic is indicated at right (Permutation test). * p<0.05, ** p<0.01, ***p<0.001

We next tracked the history of all cells to determine the origin or “source” of each splitter neuron in the preceding session. For each splitter neuron from the second included session onwards, we tracked whether that cell was a splitter neuron of any type in the preceding session or was inactive (neurons below the activity threshold or undetected by our ROI extraction algorithm). We found that previously inactive cells were the largest source category to all types of splitter neurons in 85.39% of recording sessions, and contributed an average of 55.37% of splitter neurons per session (**Figure 4b**). Turn*phase splitter neurons were the second largest category contributor to splitter neurons of all types, contributing on average 22.83% of all splitter neurons. In addition to showing the immediate integration of newly active cells into the coding population, this result suggests that representation of task variables in single units becomes less specific over time, where each cell becomes less likely to encode both task phase and turn direction.

The above result on splitter neuron sources suggests that changes in the distribution of single unit responses are, to a large degree, driven by the splitting type a newly active neuron assumes rather than transitions between different splitting types. Indeed, the proportion of splitter types of newly active cells closely matched the distribution of splitter types overall: new cells were more likely to become turn*phase splitter neurons rather than turn-only or phase-only splitter neurons (Turn*Phase vs. Turn: z=4.898, p=9.665e-07; Turn*Phase vs. Phase: z=4.804, p=1.554e-06; Wilcoxon signed-rank test) (**Figure 4c**). Additionally, the changes in this distribution of newly active cells over the course of recordings closely matched those observed for all splitter neurons (**Figure 2d**): while newly active cells on all days were more likely to be turn*phase splitter neurons than other types, this likelihood significantly decreased over time (rho=0.419, p=0.014) and the proportion of new cells allocated to turn splitter neurons on the stem significantly increased (rho=0.373, p=0.030; Spearman rank correlation), while those for phase splitter neurons and non-splitters were stable (rho=0.209, p=0.237 and rho=0.203, p=0.249 respectively) (**Figure 4d**).

Splitter and place neurons on the return arms were also found to be equally stable and primarily derived from newly active cells, but the distribution of cells newly active on the return arms among splitter types did not change over time, again suggesting the redistribution of splitter neurons is related to memory load (**Supplementary Figure 4**).

These results show that the changing distribution of single unit responses is primarily attributable to changes in the allocation of new cells to encode task variables, rather than unequal stability of different splitter types.

## Discussion

We recorded cells in dorsal CA1 of the hippocampus in mice performing a Delayed Non-Match to Place task over several sessions. In tracking the same populations of cells, we found that there was heterogeneity in the stability of task-related representations. Many single cells exhibited context-dependent modulation in their calcium activity while the animal was in the same spatial location, replicating earlier findings that demonstrate that hippocampal place cells encode the behavioral context in addition to spatial position (Griffin et al., 2007). We found that the distribution of context-dependent responses among neurons was not stable over the course of recordings: the proportion of task phase splitter neurons was stable, the proportion of turn direction splitter neurons increased, and the proportion of turn*phase splitter neurons decreased. We found this change was not attributable to variable stability of each splitter phenotype, but instead appeared to be due to how newly active cells were allocated to different splitter types. In spite of cell turnover and changes in the representation of task features among single neurons, ensemble-level population representations for different trial types were stably segregated over many recording sessions. These data demonstrate that the hippocampal representation of ongoing experience can undergo reorganization at the single neuron level while minimally impacting population level coding.

Representations may change in different ways over time during stable behavior based on competing demands on memory reorganization. Generalization emphasizes the similarities across experiences to aid in the transfer of learning across contexts, while orthogonalization makes representations more distinct to mitigate interference between contexts. Both mechanisms are important for spatial navigation and episodic memory (Hasselmo & Wyble, 1997; Kumaran & McClelland, 2012; McNaughton & Morris, 1987; Norman & O’Reilly, 2003; Schapiro, Turk-Browne, Botvinick, & Norman, 2017; Treves & Rolls, 1994; Winocur, Moscovitch, & Bontempi, 2010), and both processes are observed in fMRI studies using behavioral tasks with multiple demands (Brown and Stern 2014; Chanales et al. 2017). However, the interplay of generalization and orthogonalization in the long term reorganization of memory has not been previously studied at the single neuron level in a dynamically evolving neural circuit. Representations of different trial types may become more orthogonalized and distinct, following the precedent set by many studies on learning (Komorowski et al. 2009; McKenzie et al. 2013; Chanales et al. 2017). Alternatively, representations could become more schematic through generalization as the animals become over-trained on the task, perhaps preserving only those distinctions relevant to performing the task. At the single neuron level, we observed a result consistent with the generalization hypothesis: a decreasing number of turn*phase splitter neurons (which encode a single experience: a route to a single destination during a single task phase) and an increasing number of turn splitter neurons (which encode multiple experiences: routes to the same destination during multiple task phases). But at the population level, we instead observed a highly stable representational structure.

Studies which report orthogonalizing change in hippocampal coding properties typically examine an initial learning phase, comparing data from before and after a subject reaches a performance criterion, often in a single session (Kobayashi et al. 2003; Komorowski, et al. 2009; McKenzie et al. 2013). Because our recordings began after animals had received considerable experience with the maze environment during the pre-training phase, we may have captured a set of operational demands unlike initial learning. To reconcile our finding of generalization with previous reports of orthogonalization, we propose that both mechanisms act on the organization of memory but at different timescales: orthogonalization dominates an early, fast encoding process which emphasizes the uniqueness of current experiences, while generalization acts as a slower refinement of existing memory representations by finding statistical regularities; both of these processes likely involve regions outside the hippocampus (Ghosh & Gilboa, 2013; Koster et al., 2018; Lewis, Knoblich, & Poe, 2018). This distinction suggests that it is more appropriate for our work to be framed in terms of long-term mechanisms of memory stability, rather than those which are relevant to shaping the initial learning and encoding process.

Divergent expectations for short and long-term memory organization are apparent when comparing our results to a previous report which employed a similar task to ours in which human participants navigated partially overlapping trajectories in a virtual environment (Chanales et al., 2017). The authors found that the hippocampal voxel activity patterns for overlapping trajectory segments grew more distinct from each other over the course of learning, while patterns for non-overlapping segments did not change in their representational similarity. Our results parallel this finding in showing that conflicts between behavioral responses in overlapping locations (experienced on the central stem in the DNMP task) can drive changes in the neural representation while representations for non-overlapping segments remain stable (return arms, **Supplementary Figure 3,4**). However, unlike Chanales and colleagues, we did not observe a population-level increase in discriminability of overlapping segments, which could be explained by the fact that their study was conducted in a single session while ours ran for multiple weeks.

Prior studies have attributed a working memory role to the hippocampus in DNMP and other alternation tasks. Working memory accounts propose that on short, behaviorally relevant timescales the hippocampus maintains a representation of the previous trial to inform future behavior. This interpretation was prompted by findings that hippocampal lesions produce performance deficits in alternation tasks which involve a delay (Hampson et al. 1999; Dudchenko et al. 2000) and by correspondence between during delay period neural activity and upcoming turn directions (Deadwyler et al. 1996). However, alternation tasks cannot distinguish between prospective and retrospective coding (see Frank et al. 2000), meaning delay and central stem activity could represent a previous trial or upcoming trajectory.

We suggest instead that continued involvement of the hippocampus in distinctly representing overlapping spatial trajectories may be appropriate for self-localization within an existing spatial memory map (Redish & Touretztky, 1998). It was previously assumed that task splitter neurons reflected respective encoding and retrieval demands for Study and Test trials (Griffin et al. 2007); the self-localization interpretation suggests instead that task phase splitters instead encode immediate history of the stem traversal, whether the current trial began by being placed in the maze by the experimenter (Study) or being released from the delay area (Test). Self-localization assumes neither that the animals are sensitive to our conception of the task nor that encoding and retrieval “modes” be expressed as measurably different patterns of activity in CA1. The lack of neurons that code exclusively for Task Phase on the return arms (**Supplementary Figure 3**), where the trial-start behavioral cue is less salient, is consistent with this hypothesis. The strictest interpretation of task phase splitting as self-localization suggests it acts as a code to distinguish slightly different routes to the same reward destination (Grieves et al. 2016). Task phase splitting (**Figure 2**) and delay period splitting (Deadwyler et al. 1996) could together contribute to self-localization within a cognitive map of the task that links longer sequences of events through the maze, wherein overlapping trajectories begin on the central stem, pass down one side arm, linger in the delay area, and then pass again through the stem and onto the other side arm (Hasselmo, 2008). Task phase splitting on the central stem is similar to many other findings of context-dependent place-cell activity (Ferbinteanu & Shapiro, 2003; Frank et al., 2000; Hasselmo, 2008; Sun, Yang, Martin, & Tonegawa, 2019). Disambiguating the working-memory and self-localization accounts of splitter neuron activity will require designing tasks that use behavioral and spatial cues that are consistent across distinct but overlapping behaviors.

Our results here show that the stability of hippocampal representations is heterogeneous, displaying different rates of change in task-relevant activity across cognitive demands, maze locations, and levels of analysis. These changes are largely attributable to cells’ changes in the allocation of newly active cells among task-modulated activity types, as well as individual cells’ transitioning from coding both task dimensions to just coding for one. Together, the results suggest that reorganization of memory representations actively reshapes hippocampal memories. Future studies should seek to clarify the behavioral parameters which predict the rate of cell replacement, the allocation of newly active cells, and the cellular and network mechanisms which mediate them.

## Acknowledgments

This work was supported by the following grans: U.S. Office of Naval Research MURI N00014-16-1-2832, National Institutes of Health R01 MH052090, National Institutes of Health R01 MH 051570, and National Institutes of Health MH060013. We would like to thank Howard Eichenbaum, who initially conceived of the experimental questions and design and secured funding to perform them, but passed away in July of 2017 before the analysis had progressed significantly. We additionally thank Ian Davison for providing extensive consultation during data analysis and manuscript preparation, Jay Bladon, Dan Sheehan, and Catherine Mikkelsen for input on results, analysis and interpretation, Elin Anwar and Alex Terzibachian for help with data processing, and Dan Orlin, Wing Ning, Denise Parisi, Shelley Russek, and Sandra Grasso for administrative support. We would like to acknowledge the GENIE Program, specifically Vivek Jayaraman, PhD, Douglas S. Kim, PhD, Loren L. Looger, PhD, Karel Svoboda, PhD from the GENIE Project, Janelia Research Campus, Howard Hughes Medical Institute, for providing the GCaMP6f virus. Finally, we would like to acknowledge Inscopix, Inc. for making single-photon calcium imaging miniscopes widely available, and specifically Lara Cardy and Vardhan Dani for all their technical support throughout the experiment.

## Supplementary Data Tables

**Table 1:** Proportion comparisons splitter neurons, Wilcoxon signed-rank test, on STEM and ARMS

|  | z, STEM | p, STEM | z, ARM | p, ARMS |
| --- | --- | --- | --- | --- |
| Turn vs. Phase | 0.016379 | 0.98693 | 5.3731 | 7.7397e-08*** |
| Turn vs. Conj. | 5.2861 | 1.2497e-07*** | 2.9222 | 0.0034756** |
| Phase vs. Conj. | 5.2861 | 1.2497e-07*** | 5.3731 | 7.7397e-08*** |
| Conj. vs. Neither | 5.3586 | 8.3874e-08*** | 5.3732 | 7.7337e-08*** |
| Turn vs. Neither | 4.111 | 3.9373e-05*** | 5.3731 | 7.7397e-08*** |
| Phase vs. Neither | 4.5033 | 6.69e-06*** | 0.65823 | 0.51039 |

**Table 2:** Wilcoxon rank-sum test statistics for comparisons between population vector correlations in each spatial bin

| Spatial Bin: | VS Self vs LvR z-value | VS Self vs LvR p-value | VS Self vs SvT z-value | VS Self vs SvT p-value | LvR vs SvT z-value | LvR vs SvT p-value |
| --- | --- | --- | --- | --- | --- | --- |
| 1 | 8.312 | 9.44E-17*** | 9.653 | 4.75E-22*** | 2.924 | 0.004*** |
| 2 | 9.807 | 1.05E-22*** | 8.855 | 8.39E-19*** | -0.518 | 0.605 |
| 3 | 10.782 | 4.18E-27*** | 8.22 | 2.03E-16*** | -3.831 | 0.0001*** |
| 4 | 11.464 | 2.01E-30*** | 7.345 | 2.06E-13*** | -6.137 | 8.40E-10*** |
| 5 | 11.79 | 4.41E-32*** | 0.612 | 3.79E-11*** | -0.755 | 8.84E-15*** |
| 6 | 11.82 | 3.10E-32*** | 5.941 | 2.83E-09*** | -8.599 | 8.05E-18*** |
| 7 | 12.086 | 1.26E-33*** | 5.108 | 3.25E-07*** | -9.222 | 2.93E-20*** |
| 8 | 12.177 | 4.11E-34*** | 5.164 | 2.42E-07*** | -9.491 | 2.30E-21*** |

**Table 3:** Wilcoxon rank-sum test z and p values comparing each population vector correlation type across day lags for means of correlations in bins 1 and 2.

| Day Lag | Vs Self vs LvR z Values | Vs Self vs LvR p Values | Vs Self vs SvT z Values | Vs Self vs SvT p Values | LvR vs SvT z Values | LvR vs SvT p Values |
| --- | --- | --- | --- | --- | --- | --- |
| 1 | 8.088 | 6.09E-16*** | 7.657 | 1.90E-14*** | -0.187 | 0.852 |
| 2 | 6.893 | 5.48E-12*** | 6.062 | 1.35E-09*** | -0.614 | 0.54 |
| 3 | 5.527 | 3.25E-08*** | 4.446 | 8.77E-06*** | -1.106 | 0.269 |
| 4 | 5.29 | 1.22E-07*** | 4.721 | 2.35E-06*** | -1.467 | 0.143 |
| 5 | 5.0951 | 3.49E-07*** | 5.139 | 2.77E-07*** | -0.255 | 0.799 |
| 6 | 4.47 | 7.82E-06*** | 4.47 | 7.82E-06*** | -0.125 | 0.9 |
| 7 | 5.018 | 5.23E-07*** | 4.283 | 1.84E-05*** | -0.465 | 0.643 |
| 8 | 3.715 | 0.0002*** | 3.028 | 0.003** | -0.473 | 0.637 |
| 9 | 3.541 | 0.0004*** | 3.575 | 0.0004*** | 0.622 | 0.5341 |
| 10 | 3.152 | 0.002** | 3.727 | 0.0002*** | 0.63779 | 0.524 |
| 11 | 4.37 | 1.25E-05*** | 2.738 | 0.006** | -1.254 | 0.21 |
| 12 | 3.285 | 0.001*** | 1.749 | 0.08 | -1.556 | 0.12 |
| 13 | 2.88 | 0.004** | 2.235 | 0.026* | -0.437 | 0.663 |
| 14 | 2.67 | 0.008** | 2.203 | 0.028* | -0.302 | 0.763 |
| 15 | 2.339 | 0.019* | 1.143 | 0.253 | -0.844 | 0.399 |
| 16 | 0.855 | 0.393 | 0.479 | 0.633 | -0.5 | 0.617 |

**Table 4:**
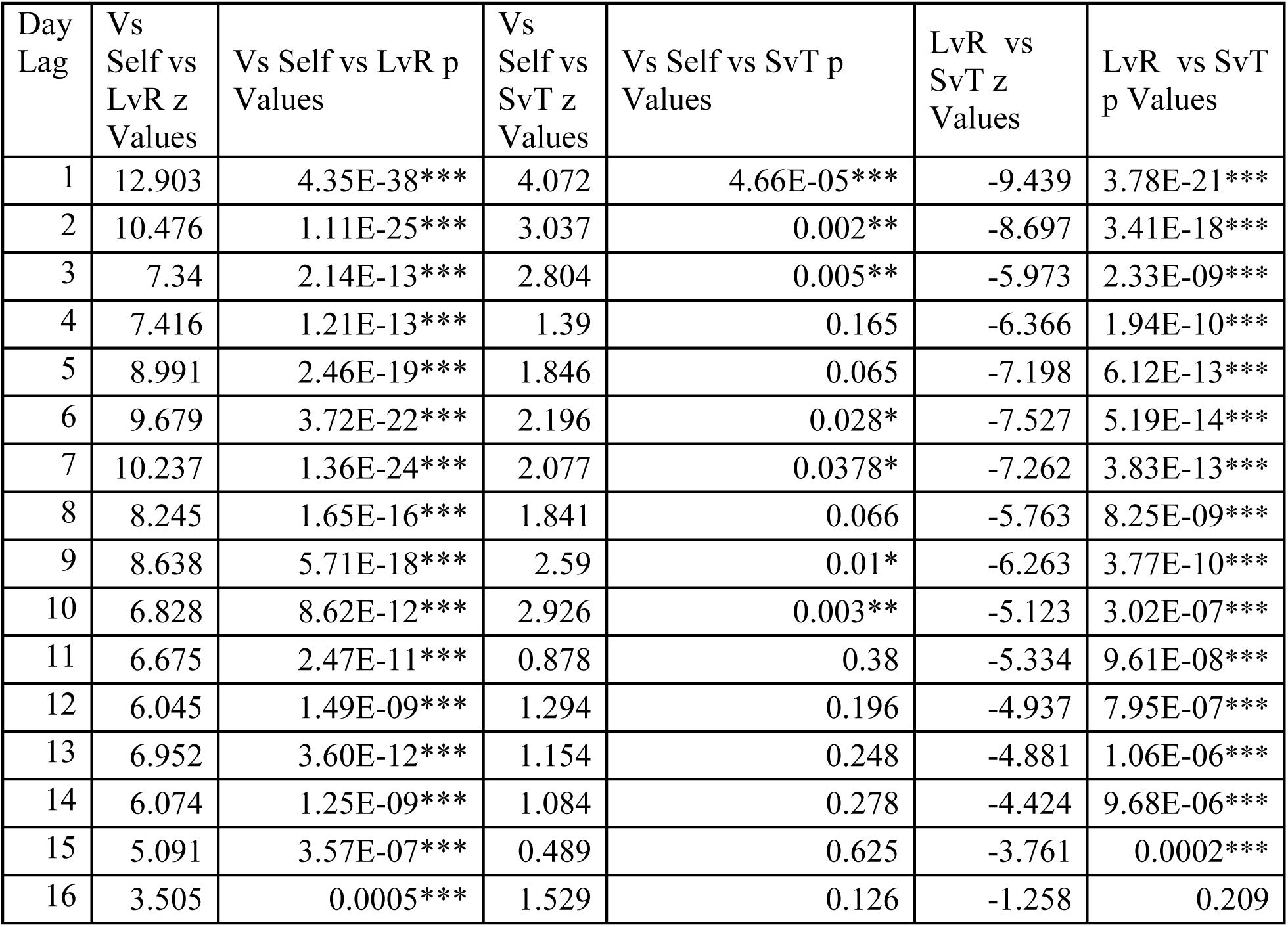
Wilcoxon rank-sum test z and p values comparing each population vector correlation type across day lags for means of correlations in bins 7 and 8.

## Software and Data availability

Software used in our analysis is freely available on GitHub. TENASPIS is available at https://github.com/SharpWave/TENASPIS, and all other analysis software is available at https://github.com/samjlevy/CaImageRelated. Data can be made available from the authors upon reasonable request.

---

**Supplementary Figure 1.**
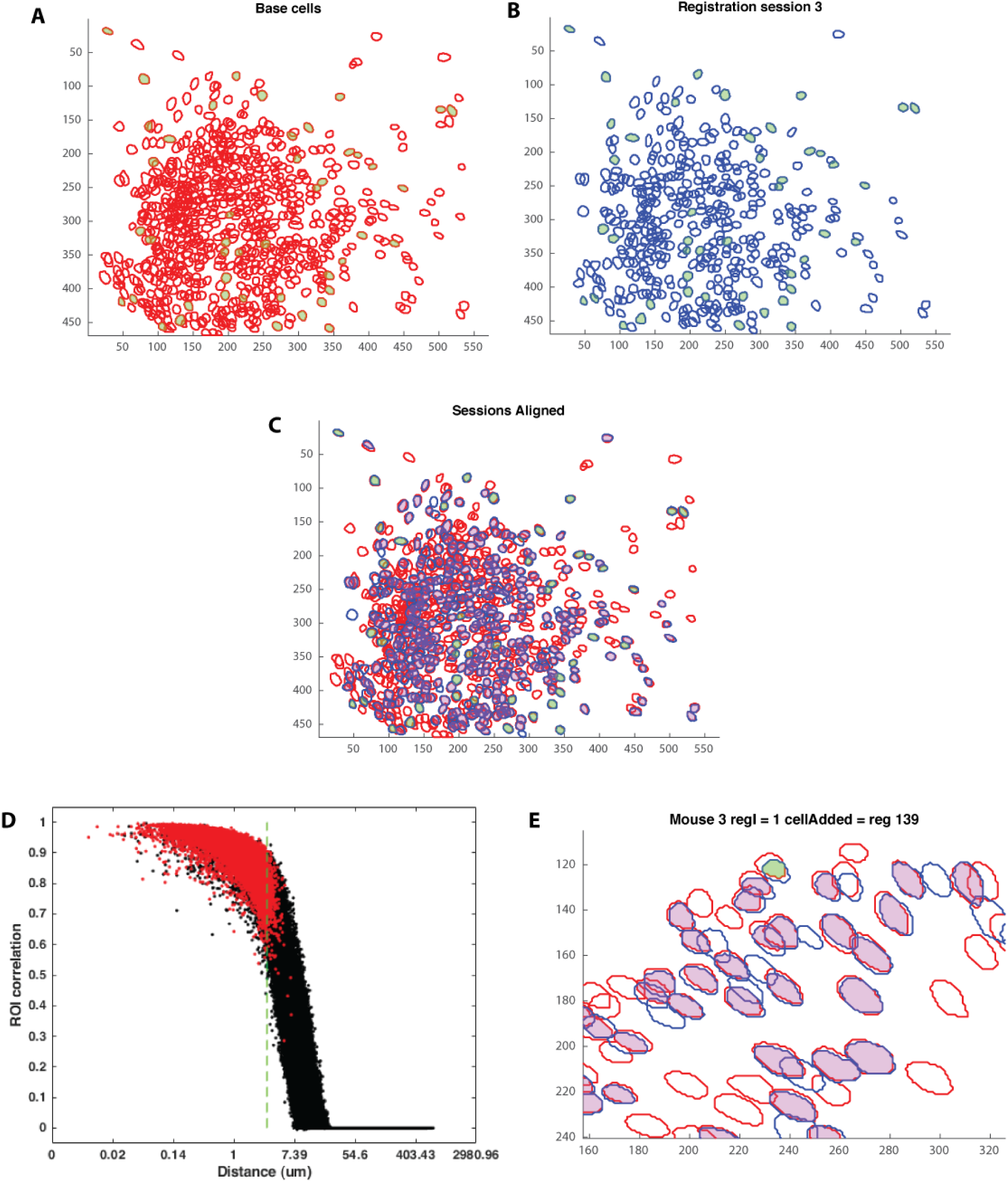
***A,B***Cell ROI outlines for the base session (**A**) and one registered session (**B**) for one mouse. Green filled-in cells are manually selected “anchor cells” used to compute the affine transformation for alignment. ***C***, Overlaid base session in red and registered session in blue, same as ***A,B***. “Anchor cells” filled in green, and other registered cells are filled in purple. ***D,*** Scatter plot showing relationship between ROI correlation and center-to-center distance for every pair of cells in each base-registered session pair. Registered cells are marked in red. Green dashed line indicates 3 um threshold used during registration. X-axis is log-scaled. ***E,*** Enlarged section of a registered session from a different mouse from ***A-C*** illustrating a manually registered cell (filled in green). This cell was skipped by the algorithm because the centers in the base and registered sessions were further apart than the 3um threshold (3.316um, ROI correlation 0.757). This cell was added manually based on its relative alignment to other cells successfully registered and the similarity of ROI outlines.

**Supplementary Figure 2.**
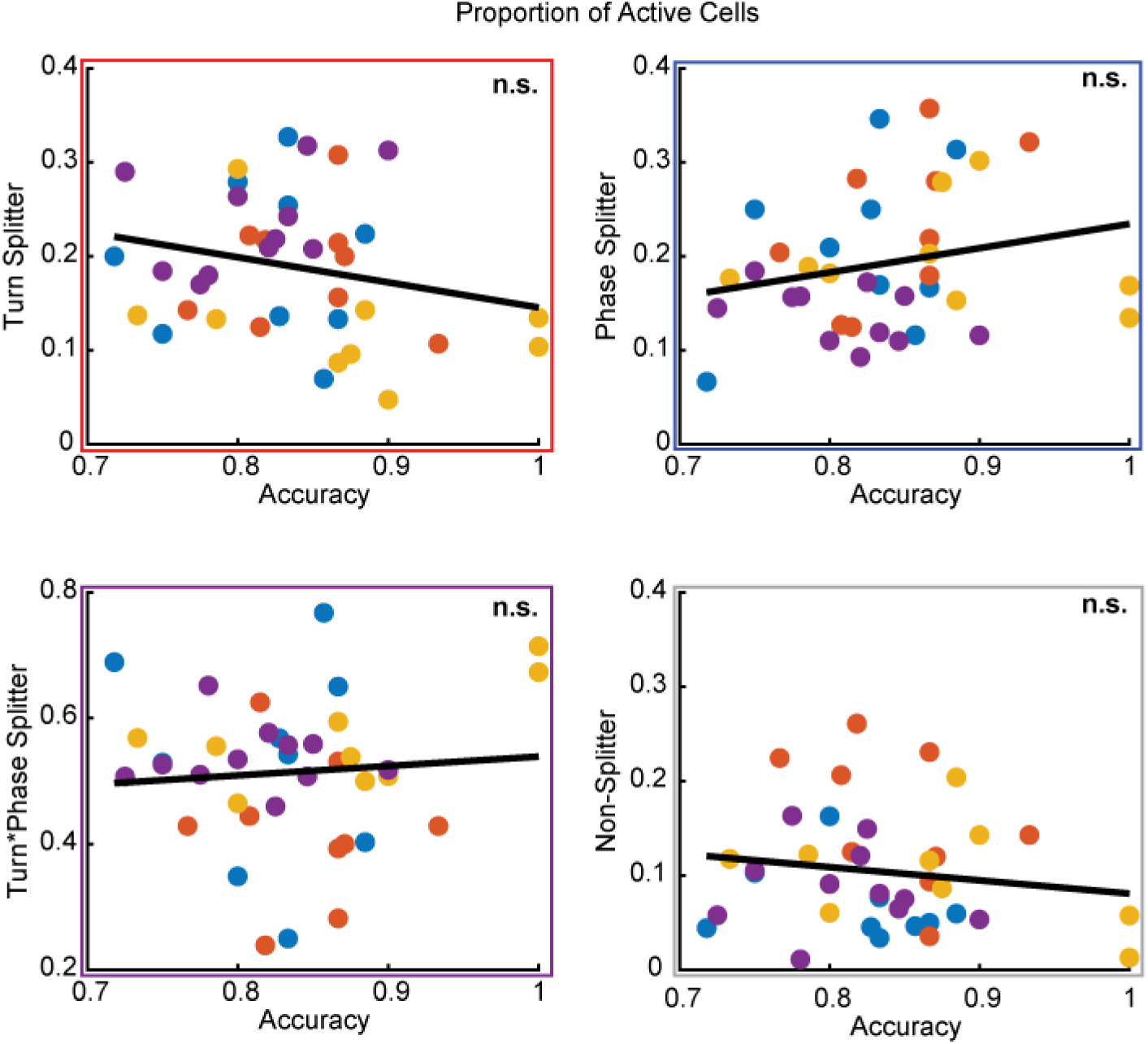
Correlation between proportions of splitter cells out of total active cells with animal’s performance in that session. Dot color refers to each mouse, each point is a single session. Black line is best fit linear regression. Box color indicates splitter type detailed in y-axis. Significance is calculated with a spearman rank correlation between the proportion of splitter cells and session accuracy. Turn rho=-0.210, p=0.206. Phase rho=0.217, p=0.190. Turn*Phase rho=-0.030, p=0.857. Non-splitter rho=-0.136, p=0.417. * p<0.05, ** p<0.01, ***p<0.001

**Supplementary Figure 3.**
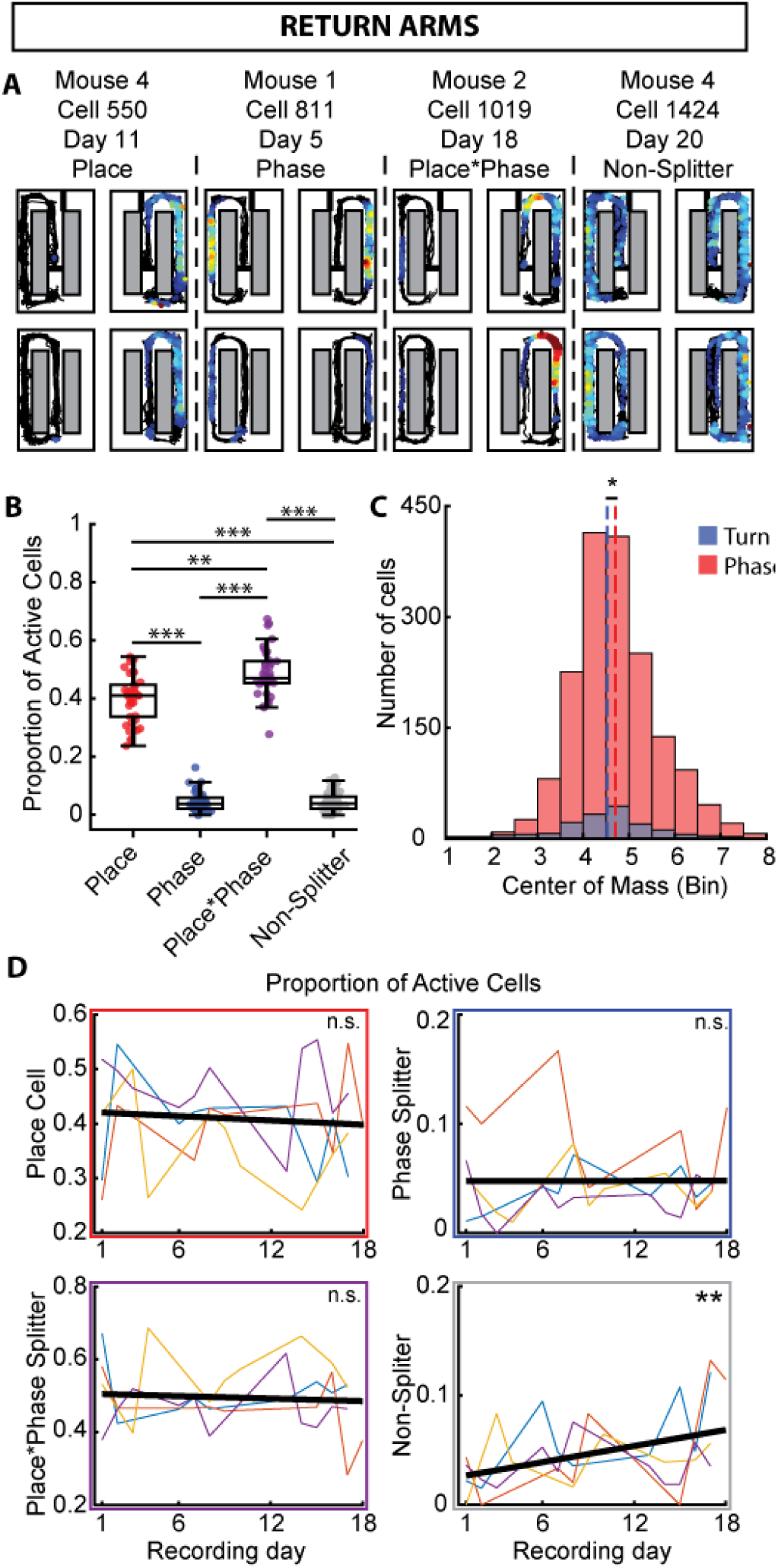
***A,*** Example activity maps for each type of splitter on the return arms. Warmer colors indicate higher transient likelihood. ***B,*** Proportions of splitter cells out of the total active cell population on each day for all animals. Box shows inter-quartile range and middle line shows median. Statistic: Wilcoxon signed-rank test. ***C,*** Distribution of centers-of-mass of event activity for Turn and Phase splitter neurons. Statistic: Mann-Whitney U-test. ***D,*** Proportion of splitter neurons in individual animals (unique colors) and group regression (black) over the course of the experiment. Significance indicates between all included recording sessions (n=38). Statistic: Spearman rank correlation (Proportion of splitters by recording day number). *p<0.05, **p<0.01, ***p<0.001

**Supplementary Figure 4.**
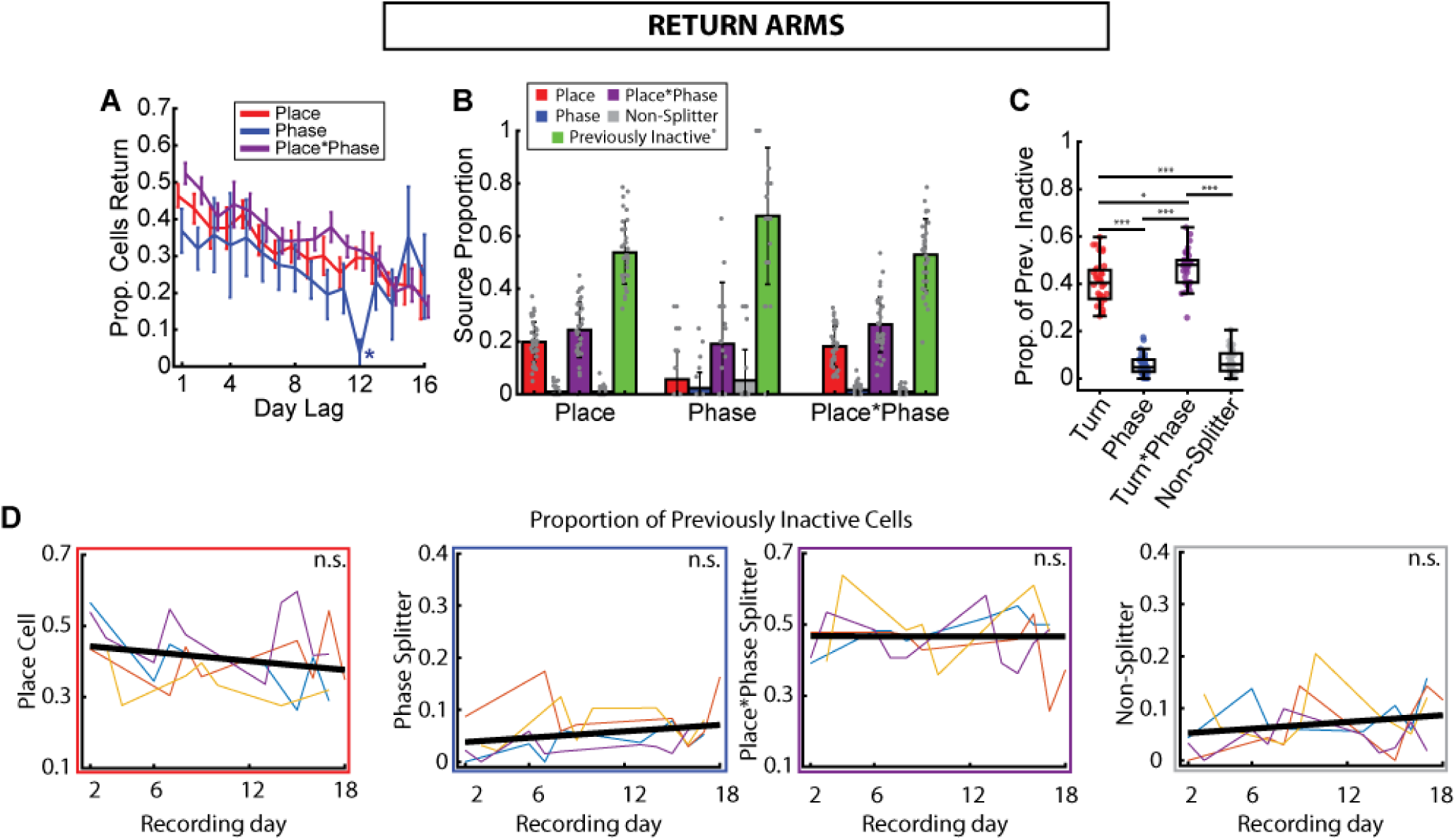
***A,*** Proportion of cells that are still present at increasing day lags. Statistic: Wilcoxon signed-rank test. ***B,*** Proportion of each splitter type by what that cell was on the prior day of recording. ***C,*** Proportion of each splitting phenotype among each recording day’s set of previously inactive cells (from second recording day forward). Statistic: Wilcoxon signed-rank test. ***D,*** Changes in the distribution of splitting phenotypes among previously inactive over the course of recordings. Colored lines are individual animals, black line is best fit regression. Statistic is indicated at right (Permutation test). * p<0.05, ** p<0.01, ***p<0.001

